# Background selection as baseline for nucleotide variation across the *Drosophila* genome

**DOI:** 10.1101/005017

**Authors:** Josep M Comeron

## Abstract

The constant removal of deleterious mutations by natural selection causes a reduction in neutral diversity and efficacy of selection at genetically linked sites (a process called Background Selection, BGS). Population genetic studies, however, often ignore BGS effects when investigating demographic events or the presence of other types of selection. To obtain a more realistic evolutionary expectation that incorporates the unavoidable consequences of deleterious mutations, we generated high-resolution landscapes of variation across the *Drosophila melanogaster* genome under a BGS scenario independent of polymorphism data. We find that BGS plays a significant role in shaping levels of variation across the entire genome, including long introns and intergenic regions distant from annotated genes. We also find that a very large percentage of the observed variation in diversity across autosomes can be explained by BGS alone, up to 70% across individual chromosome arms, thus indicating that BGS predictions can be used as baseline to infer additional types of selection and demographic events. This approach allows detecting several outlier regions with signal of recent adaptive events and selective sweeps. The use of a BGS baseline, however, is particularly appropriate to investigate the presence of balancing selection and our study exposes numerous genomic regions with the predicted signature of higher polymorphism than expected when a BGS context is taken into account. Importantly, we show that these conclusions are robust to the mutation and selection parameters of the BGS model. Finally, analyses of protein evolution together with previous comparisons of genetic maps between *Drosophila* species, suggest temporally variable recombination landscapes and thus, local BGS effects that may differ between extant and past phases. Because genome-wide BGS and temporal changes in linkage effects can skew approaches to estimate demographic and selective events, future analyses should incorporate BGS predictions and capture local recombination variation across genomes and along lineages.

## AUTHOR SUMMARY

The removal of deleterious mutations from natural populations has potential consequences on patterns of variation across genomes. Population genetic analyses, however, often assume that such effects are negligible across recombining regions of species like *Drosophila*. We use simple models of purifying selection and current knowledge of recombination rates and gene distribution across the genome to obtain a baseline of variation predicted by the constant input and removal of deleterious mutations. We find that purifying selection alone can explain a major fraction of the observed variance in nucleotide diversity across the genome. The use of this baseline of variation as null expectation also exposes genomic regions under other selective regimes, including more regions showing the signature of balancing selection than would be evident when using traditional approaches. Our study also shows that most, if not all, nucleotides of the *D. melanogaster* genome are influenced by nearby deleterious mutations and thus the frequent assumption that sites evolve independently of one another is likely unwarranted across the entire genome. Additionally, the study of rates of protein evolution suggests that the recombination landscape across the genome has changed in the recent history of *D.melanogaster*, a factor that can skew analyses designed to estimate the strength and frequency of adaptive events. Together, these results illustrate the need for new and more realistic theoretical and modeling approaches to capture demographic and selective events.

## INTRODUCTION

The causes of the variation observed within natural populations have been a long-standing question in evolutionary and genetic studies. Particular insight into these causes can be gained by analyzing the distribution of nucleotide diversity across genomes, where species- and population-specific parameters such as the number of individuals, environmental factors, or demography are constant. A number of population genetics models have been put forward to explain this intra-genomic variation in diversity, often including the predicted consequences that selection acting at a genomic site impinges on genetically linked sites, either neutral or under selection themselves (i.e., models of ‘selection at linked sites’; [1–4] and references therein). Although there is general agreement that selection at linked sites can play a role shaping levels of variation, there is still intense debate and research on the selective nature of the mutations causing such effects (e.g., beneficial or deleterious) and whether the same causes can be applied to different species [4–6].

Strongly beneficial mutations rise rapidly to fixation and hitchhike adjacent linked sites, causing a fingerprint of reduced intra-specific variation around the selected site known as a ‘selective sweep’ (the HHss model; [2,3,7–11]). A qualitatively similar outcome can be generated by another model of selection and hitchhiking effects at linked sites without requiring adaptive changes, just as a result of the continuous input of strongly deleterious mutations and their removal by natural selection (the background selection (BGS) model [1,4,12–16]). Both models also predict that the consequences of selection removing adjacent diversity diminish when genetic recombination increases, a general pattern that has been observed in many species when comparing genomic regions with high and low (or zero) recombination rates (reviewed in [1,3–5]).

The magnitude and distribution of recombination rates across genomes play key roles in predicting the consequences of selection on adjacent variation. In humans, for instance, the presence of large recombination cold spots raised the possibility that BGS could reduce polymorphism levels at specific genomic regions. In agreement, recent analyses using models of purifying selection rather than purely neutral ones suggest that patterns of nucleotide diversity across the human genome are consistent with BGS predictions [17–21]. In the model system *D. melanogaster*, low-resolution recombination maps described limited or absent recombination near sub-telomeric and –centromeric regions whereas recombination outside these sub-telomeric and –centromeric regions (i.e., across *trimmed* chromosome arms) has been often assumed to be both high and homogeneously distributed. As a consequence, variation in nucleotide diversity across trimmed chromosome arms has been mostly attributed to positive selection and selective sweeps ([4,5,22–29]; but see [30]).

There are, however, several reasons to believe that BGS effects could be significant in *D.melanogaster* as well. First, compared with humans, *D. melanogaster* has a more compact genome and a larger effective population size (*N_e_*), predicting tighter genetic linkage between genes and stronger purifying selection, respectively, and both factors forecast greater BGS effects. Second, recent whole-genome studies of recombination rates in *D. melanogaster* exposed extensive heterogeneity in the distribution of crossover rates even after removing sub-telomeric and centromeric regions [31]. This high degree of variation in recombination rates across *D. melanogaster* chromosomes is observed when recombination is obtained from a single cross of two specific strains [31,32] as well as when analyzing a species’ average obtained from combining genetic maps from crosses of different natural strains [31]. The presence of coldspots of recombination embedded in chromosomal regions assumed to have high recombination rates, therefore, provides the opportunity for BGS to play a more significant role across broader genomic regions than previously anticipated [31]. Finally, Charlesworth [33] has recently showed that BGS effects are predicted to be detectable in the middle of recombining chromosome arms in *D. melanogaster.*

The consequences of BGS at a given nucleotide position in the genome (focal point) can be described by the predicted level of neutral nucleotide diversity when selection at linked sites is allowed (π) relative to the level of diversity under complete neutrality and free recombination between sites (π_0_), with *B* = π/π_0_ [12–15]. Therefore, *B* ∼ 1 would indicate negligible BGS effects whereas *B* << 1 would suggest very strong BGS and a substantial reduction in levels of neutral diversity. *B* can also be understood in terms of a reduction in *N_e_* and therefore variation in *B* forecasts differences in levels of diversity within species but also differences in the efficacy of selection, which can be approximated by the product of *N_e_* and the selection coefficient *s*. Note, however, that the prediction about reduced efficacy of selection is a qualitative one since there is no simple scalar transformation of *N_e_* influenced by selection at linked sites that allows estimating probabilities of fixation of selected mutations [34,35]. Thus, a comprehensive study of the predictive power of BGS to explain natural variation across genomes needs to show that, 1) conditions exists across a genome to generate significant overall effects reducing *B,* 2) *B* varies across the genome, and 3) regions with reduced *B* are associated with reduced levels of polymorphism and efficacy of selection (e.g., detectable on rates of protein evolution).

Here, we investigated what is the fraction of the *D. melanogaster* genome that is influenced by BGS and how much of the observed variance in patterns of intra-specific variation and rates of evolution across this genome can be explained by BGS alone. Importantly, to obtain a sensible BGS baseline that could be used to test for positive selection and other departures from neutrality, we investigated BGS models that are purposely simple and independent of nucleotide variation data. Additionally, we studied whether our conclusions are sensitive to parameters of the BGS model. To this end, we expanded approaches previously applied to investigate human diversity [17,18] to now estimate BGS effects across the *D. melanogaster* genome.

In all, we generated a detailed description of the consequences of purifying selection on linked sites at every 1kb along *D. melanogaster* chromosomes under several BGS models. Our results show that BGS likely plays a detectable role across the entire genome and that purifying selection alone can explain a very large fraction of the observed patterns of nucleotide diversity in this species. Notably, we show that these conclusions are robust to different parameters in the BGS models. The use of a BGS baseline also uncovers the presence of regions with the signature of a recent selective sweep and, less expected, numerous instances of balancing selection. Furthermore, analyses of rates of protein evolution suggest that the recombination landscape has changed recently along the *D. melanogaster* lineage thus generating disparity between short- and long-term *N_e_* at many genomic positions. We discuss the advantages of incorporating BGS predictions across chromosomes and the potential consequences of temporal variation in recombination landscapes when estimating demographic and selective events.

## RESULTS

### The BGS Models

BGS expectations (i.e., estimates of *B*) were obtained for every 1-kb region across the whole genome as the cumulative effects caused by deleterious mutations at any other site along the same chromosome (see **Materials and Methods** for details). These estimates of *B* were based on BGS models that include our current knowledge of genome annotation at every nucleotide site of the genome and high-resolution recombination landscapes in *D. melanogaster* that distinguish between crossover and gene conversion rates [31]. These models also incorporate the possibility that strongly deleterious mutations occur at sites that alter amino acid composition as well as at a fraction of sites in noncoding sequences. The inclusion of deleterious mutations in noncoding sequences allows taking into account the existence of regulatory and other non-translated functional sequences, either in introns and 5’- and 3’- flanking UTRs, or in intergenic regions [22,33,36–38]. For each category of selected sites (nonsynonymous, intronic, UTR, or intergenic) we used the proportion of constrained sites (*cs*) estimated for *D. melanogaster* [22,37,38] as the fraction of sites with deleterious fitness consequences when mutated [33]. In terms of recombination rates, we studied BGS predictions following the standard approach of including crossover as the sole source of recombination (hereafter models M_CO_) and also when combining the effects of crossover and gene conversion events (models M_CO+GC_) to better quantify the true degree of linkage between sites in natural populations.

The distribution of deleterious fitness effects (DDFE) and the diploid rate of deleterious mutations per generation (*U*) are parameters that have direct implications on estimates of *B* but are more difficult to establish experimentally. Although a gamma distribution has been proposed a number of times for deleterious mutations [39–44], a log-normal DDFE allows capturing the existence of lethal mutations and fits better *D. melanogaster* polymorphism data [45,46]. Additionally, a log-normal DDFE predicts a higher fraction of mutations with minimal consequences removing linked variation than a gamma DDFE and, ultimately, weaker BGS effects (see **Materials and Methods).** Therefore, the use of a log-normal DDFE can be taken as a conservative approach when inferring the magnitude of BGS effects.

Direct estimations of deleterious mutation rates are still fairly limited. In *D. melanogaster,* initial analyses of mutation accumulation lines estimated a mutation rate for point mutations and small indels (*u*) of ∼ 8.4 × 10^−9^ /bp /generation and a diploid rate of deleterious mutations per generation (*U*) of ∼ 1.2 [47]. Nevertheless, one of the lines used in this study had an unusually high mutation rate [48] and more recent studies suggest *u* ∼ 4-5 × 10^−9^ (*U* ∼ 0.6) for point mutations and small indels [48–50]. These lower estimates, however, do not include the deleterious consequences of transposable element (TE) insertions or the possible presence in natural populations of genotypes with high mutation rates. In fact, TEs are very abundant in natural populations of *D. melanogaster* [51–60] and have been proposed to be an important source of BGS in this species [30]. Therefore, *U* ∼ 0.6 represents a lower boundary for the deleterious mutation rate when inferring the consequences of BGS. To include the consequences of TE insertion in our BGS models, we obtained an approximate diploid insertion rate of *U*_TE_ ≥ 0.6 based on a detailed description of TE distribution in *D. melanogaster* [60] and mutation-selection balance predictions (see **Materials and Methods** for details). Thus, a genome-wide diploid deleterious mutation rate of ∼ 1.2 per generation is a reasonable approximation that captures the consequences of point mutations, small indels and the insertion of transposable elements.

To assess how robust our results and conclusions are to the parameters of the BGS model, we obtained genome-wide landscapes for *B* under eight different models, with DDFE following a log-normal or a gamma distribution (models M_LN_ and M_G_, respectively), with deleterious mutations rates that include or not TE insertions (models M_StdMut_ and M_LowMut_, respectively), and with recombination taking into account crossover and gene conversion events or only crossovers (models M_CO+GC_ and M_CO_, respectively). Unless specifically noted, we report results based on the BGS model that is most consistent with our current knowledge of gene distribution across the genome, a log-normal DDFE, a genome-wide diploid deleterious mutation rate of *U* = 1.2, and recombination rates that include crossover and gene conversion events (i.e., our default model is M_LN,StdMut,CO+GC_). We summarize the results from the BGS models in Table S1 and provide the full distribution of *B* estimates across all chromosomes in Table S2.

### Patterns of BGS across the *D. melanogaster* genome

Genome-wide estimates of *B* show a median of 0.591 and indicate that the predicted influence of BGS across the *D. melanogaster* genome would reduce the overall *N_e_* substantially relative to levels predicted by evolutionary models with free recombination (see Figure 1A). The study of the distribution of *B* along chromosomes shows that the reduction in neutral diversity is severe in a large fraction of the genome, with 19% of all 1-kb regions with *B* < 0.25 and a lower 90% CI for *B* of 0.005 (Figure 1B and Figure 2). Importantly, the distribution of *B* across *trimmed* chromosomes is also highly heterogeneous. As expected, estimates of *B* are strongly influenced by variation in local crossover rates (*c*), with a Spearman’s rank correlation coefficients (*ρ*) between *B* and *c* of 0.792 for trimmed chromosomes. As shown in Figure 3, however, there is detectable variance in *B* for a given local *c* that exposes the additional effects of long-range distribution of recombination rates and genes when estimating *B* at a focal point.

**Figure 1.**
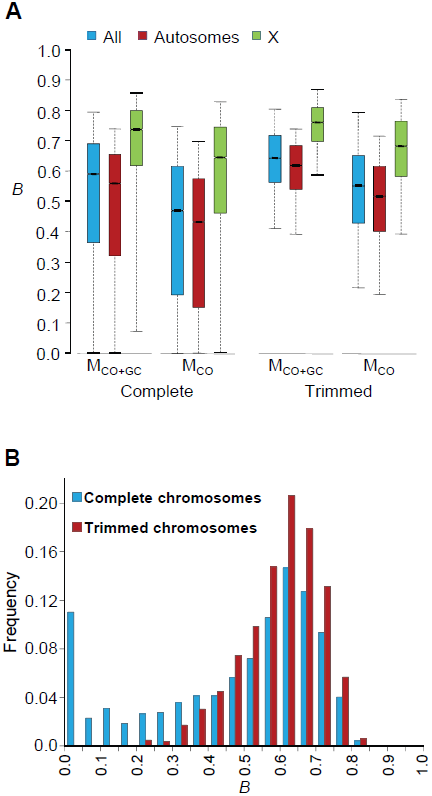
Genome-wide estimates of BGS. **(A)** Boxplots of estimates of *B* for complete or trimmed chromosomes and for models M_LN,StdMut,CO+GC_ (M_CO+GC_) and M_LN,StdMut,CO_ (M_CO_). Results shown for the complete genome, and autosomes and the X chromosome separately. The median is identified by the line inside the box, the length of the box and whiskers indicate 50% and 90% CI, respectively. **(B)** Frequency distribution of *B* estimates for complete or trimmed chromosomes under model M_LN,StdMut,CO+GC_. All results based on the analysis of 1-kb non-overlapping regions.

**Figure 2.**
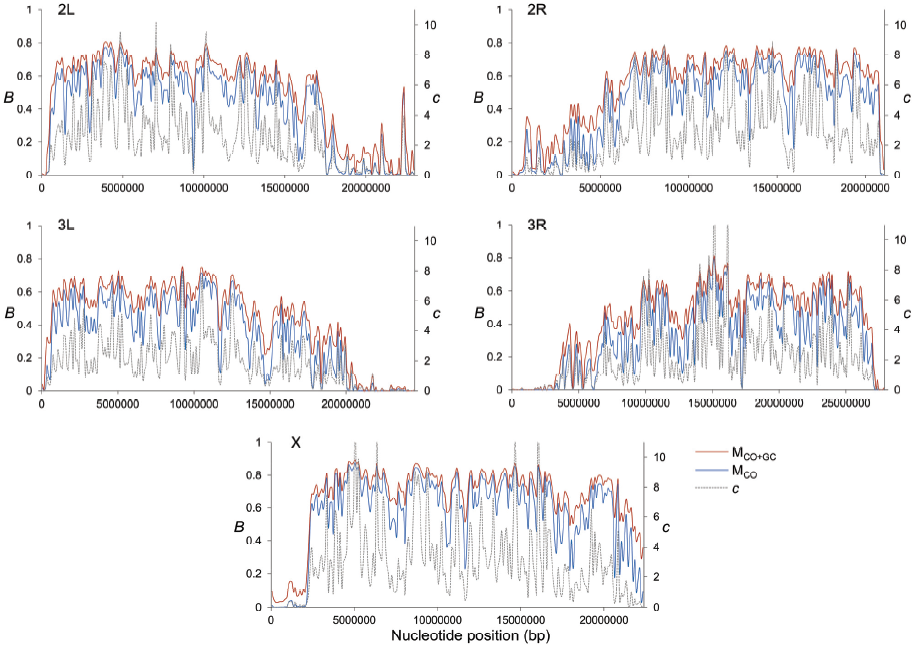
High-resolution distribution of BGS effects across the *D. melanogaster* genome. Estimates of BGS effects are measured as *B* and shown along each chromosome arm for 100-kb adjacent windows. Red and blue lines depict estimates of *B* based on models M_LN,StdMut,CO+GC_ (M_CO+GC_) and M_LN,StdMut,CO_ (M_CO_), respectively. Grey dashed lines show the distribution of crossover rates (*c*), measured as centimorgans (cM) per megabase (Mb) per female meiosis (see [31] for details).

**Figure 3.**
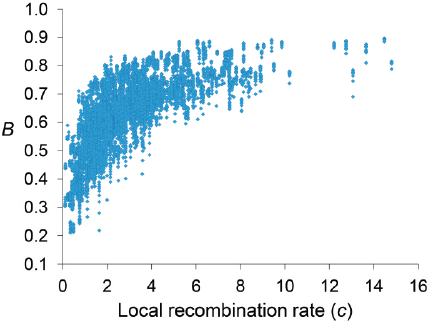
Relationship between local recombination rates and estimates of *B* across trimmed chromosomes. Local recombination rates (*c*) measured as cM/Mb per female meiosis, and estimates of *B* based on model M_LN,StdMut,CO+GC_. Results shown for 10-kb non-overlapping regions.

Median *B* across trimmed chromosome arms is 0.643, with a minimum estimate of 0.19. These results indicate that significant and variable BGS effects are expected in *D. melanogaster* not only due to sub-telomeric and -centromeric regions but also across trimmed chromosomes (see also [33]). This general conclusion does not vary qualitatively when considering BGS models with other parameters (Table S1). As expected, a model with a DDFE following a gamma distribution (M_G_) predicts stronger BGS effects and lower estimates of *B* across the genome than when the DDFE follows a log-normal (as in our default model). Under model M_G,CO+GC_ the median *B* is 0.428 and the lower 90% CI for *B* is 0.001 (median *B* across trimmed chromosomes is 0.493, with a minimum estimate of 0.007). Also anticipated, models with a lower deleterious mutation rate (models M_LowMut_) generate higher estimates of *B* than when TE insertions are taken into account (models M_StdMut_). For instance, median *B* increases from 0.591(M_LN,StdMut,CO+GC_) to 0.769 (M_LN,LowMut,CO+GC_), and from 0.428 (M_G,StdMut,CO+GC_) to 0.654 (M_G,LowMut,CO+GC_). In addition, the comparison of predictions under models with and without gene conversion shows that the standard approach of considering crossover as the only source of recombination between sites would overestimate linkage effects. Median estimates of *B* are 20 and 21% lower for models M_LN,CO_ and M_G,CO_ than for M_LN,CO+GC_ and M_G,CO+GC_, respectively. The use of only crossover rates in BGS models skews estimates of *B* particularly in regions with intermediate rates (∼0.2-2 cM/Mb), mostly across trimmed chromosomes. Both crossover and gene conversion data, therefore, need to be considered to obtain accurate estimates of the consequences of selection on linked sites and, in this case, the magnitude of BGS effects.

Finally, it is worth noting that although the different BGS models predict different point estimates and ranges of *B* across the chromosomes, all models generate *B* estimates across the genome that have virtually the same relative ranking (i.e., monotonically related; Table S3). Pairwise Spearman’s *ρ* between estimates of *B* from different BGS models range between *ρ* = 0.9856 (comparing the two most distinct models M_LN,StdMut,CO_ and M_G,LowMut,CO+GC_) and *ρ* > 0.9999 (for the four comparisons between models differing in the deleterious mutation rate).

#### No BGS-free regions in the *D. melanogaster* genome

The upper end of the distribution of *B* across the genome is also informative and relevant for population genetic analyses of selection and demography that benefit from using regions evolving not only under neutrality but also free of linkage effects. The upper 90% CI for *B* is 0.80 across the whole genome and 0.814 across the trimmed genome (Figures 1 and 2 and Table S1). Out of 119,027 1-kb regions investigated across the genome, the highest *B* is 0.897 and is located in a genomic region with very low density of genes and high crossover rate: position 5.027 Mb of the X chromosome, in the middle of a large 50-kb region with a single and short CDS and a crossover rate of *c* ∼14 cM/Mb. The 1-kb autosomal region with the least BGS effects shows *B* = 0.822. BGS, therefore, plays a significant role across the entire genome, including long introns and intergenic sequences distant from genes in the middle of regions with high recombination rates. This conclusion is unlikely to be influenced by the parameters of the BGS model. As indicated, M_G_ models predict stronger BGS effects and lower *B* and, accordingly, we observe an upper 90% CI for *B* of 0.707 and a maximum estimate of 0.87. On the other hand, models with lower deleterious mutation rates generate higher *B,* but even these models predict detectable BGS across the whole genome. M_LowMut_ models generate upper 90% CIs for *B* ranging between 0.811(M_G,LowMut,CO_) and 0.895(M_LN,LowMut,CO+GC_), and the highest 1-kb estimate of *B* obtained by any of our M_LowMut_ models is 0.947 (see Table S1 **and Figure S1)**. That is, these results strongly suggest that while there may be a fraction of sites free of selective constraints across the *D. melanogaster* genome, all sites might be, nonetheless, under the influence of selection at linked sites and BGS in particular.

#### Autosomes show stronger BGS effects than the X chromosome

The X chromosome in *D.melanogaster* recombines at a higher rate than autosomes, caused by a higher median crossover rate per female meiosis (2.48 *vs*. 1.74 cM/Mb for X and autosomes, respectively), and the fact that the X chromosome spends less time than autosomes in males (that do not recombine during meiosis). This higher recombination, in turn, forecasts weaker BGS in the X [30,61]. In agreement, Charlesworth [33] predicted weaker BGS effects (higher *B*) in the middle of the X chromosome than in the middle of an autosome under a number of models. Our study shows the same trend (see Figure 1A and Table S1), with median *B* of 0.559 and 0.736 for autosomes and the X chromosome, respectively (0.619 and 0.761 for trimmed autosomes and the X chromosome, respectively). The direct comparison of all 1-kb regions reveals that this higher *B* in the X chromosome relative to autosomes is highly significant (Mann-Whitney *U* Test, *P* < 1×10^−12^ for complete and trimmed chromosomes), with all BGS models generating the same trend and level of significance.

The ratio of observed neutral diversity for the X and autosomes (X/A) predicted by our default BGS model is 0.99 and 0.92 for complete and trimmed chromosomes, respectively. Therefore, differences in effective BGS effects at the X and autosomes can, at least in part, explain the observation that the X/A ratio of neutral diversity in several *D. melanogaster* populations is higher than the 0.75 predicted by most neutral models and a 1:1 sex ratio (see [33]). Note that X/A ratios would be overestimated if gene conversion events were not taken into account, with X/A ratios of 1.12 and 0.99 for complete and trimmed chromosomes, respectively.

### The genomic units of BGS in *D. melanogaster*

In this study, all sites along a chromosome were allowed to potentially play a role adding up BGS effects at any focal region of the same chromosome. To investigate the size of the genomic region causing detectable BGS effects in *D. melanogaster*, we estimated the size of the region surrounding a focal 1-kb needed to generate 90% of the total BGS effect obtained when considering the complete chromosome (D*_B90_* in either genetic or physical units). Equivalently, we also obtained D*_B75_* and D*_B50_* as the size of the genomic region needed to generate 75 and 50%, respectively, of the total BGS effect obtained when considering the whole chromosome.

The study of complete chromosomes shows a median genetic D*_B90_*, D*_B75_* and D*_B50_* of 5.5, 1.2 and 0.15 cM, respectively. In terms of physical distance, median D*_B90_*, D*_B75_* and D*_B50_* are 2024, 477 and 76 kb, respectively (Figure 4A). Although the overall effects of BGS are reduced along trimmed chromosomes compared to whole chromosomes, the size of the region playing a significant role in the final magnitude of *B* at a focal point is fairly equivalent, with 6.9, 1.8 and 0.21 cM for D*_B90_*, D*_B75_* and D*_B50_*, respectively (2,412, 640 and 84 kb for D*_B90_*, D*_B75_* and D*_B50_*, respectively). This genetic and physical scale, moreover, increases with crossover rates (Figure 4B). This analysis, therefore, suggests that the extent of BGS at most genes and intergenic sites across the *D. melanogaster* genome is influenced by the cumulative effects of many sites and include numerous other genes. Thus, accurate estimates of *B* in *D. melanogaster* require the study of genomic regions at the cM or Mb scale, ideally full chromosomes. Otherwise, BGS can be severely underestimated, and inferences about demographic events or other types of selection may be unwarranted. All these results are also in agreement with the previous observation that all intergenic sequences and introns across the genome are predicted to be influenced by BGS.

**Figure 4.**
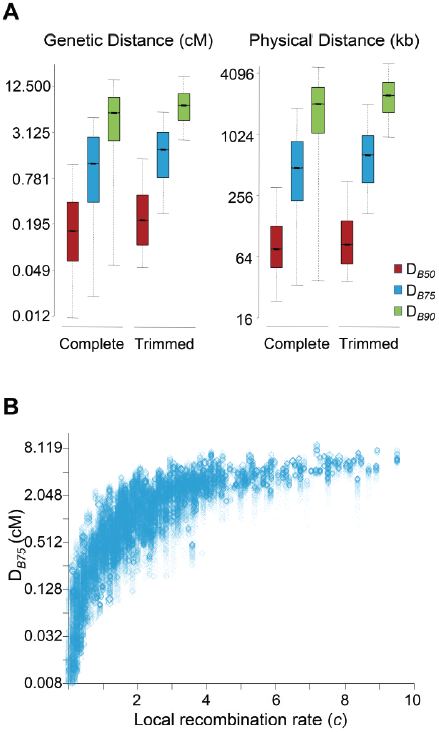
Genomic distance influencing patterns of BGS in *D.melanogaster*. **(A)** Boxplots of estimates of D*_B50_*, D*_B75_* and D*_B90_*. D*_B90_* is defined as the size of the genomic region around a focal point needed to generate 90% of the total BGS effect obtained when considering the whole chromosome. Equivalently, D*_B75_* and D*_B50_* indicate genomic distances needed to generate 75 and 50%, respectively, of the total BGS effect obtained when considering the complete chromosome. The units of D*_B_* are genetic distances (cM/female meiosis) or physical distances (kb). (See Figure 1 legend for further explanation of boxplots.) Results shown for the default model M_LN,StdMut,CO+GC._ **(B)** Relationship between local recombination rates (*c*; cM/Mb per female meiosis) and estimates of D*_B75_* in genetic distance. Results shown based on the analysis of 1-kb non-overlapping regions across the whole genome (Spearman’s ρ = 0.907, *P* < 1×10^−12^).

### Estimates of *B* are a very strong predictor of nucleotide diversity across the whole *D. melanogaster* genome

A second goal of this study was to estimate how much of the observed levels of neutral diversity across the *D. melanogaster* genome can be explained by a BGS landscape obtained independently from variation data. A strong positive correlation would not only indicate that BGS should not be ignored in population genetic analyses but also that our estimates of *B* are likely suitable as baseline to infer additional types of selection and/or demographic events. Because our best experimentally-obtained whole-genome recombination maps for crossover and gene conversion have a maximum resolution and accuracy at the scale of 100-kb [31], the predictive nature of the *B* baseline was first investigated at this physical scale.

To obtain levels of neutral diversity across the *D. melanogaster* genome, nucleotide diversity per bp (π_sil_) at introns and intergenic sequences was estimated from a sub-Saharan African population (Rwanda RG population [62]; see **Materials and Methods** for details). The comparison of estimates of *B* generated by our BGS models and levels of π_sil_ across the genome reveals a strikingly positive association (Table 1 and Figure 5). For autosomes, the correlation between *B* and π_sil_ is *ρ* = 0.770 (965 non-overlapping 100-kb regions, *P* < 1×10^−12^), and increases up to *ρ* = 0.836 (*P* < 1×10^−12^) along individual autosome arms. Equivalent results are obtained when silent diversity is analyzed separately at intergenic and intronic sites, with *ρ* = 0.736 between *B* and π_intergenic_, and *ρ* = 0.741 between *B* and π_intron_ (*P* < 1×10^−12^ in both cases). The study of individual autosome arms shows a positive association up to *ρ* = 0.799 and 0.800 for intergenic and intronic sites, respectively (*P* < 1×10^−12^ in both cases).

**Figure 5.**
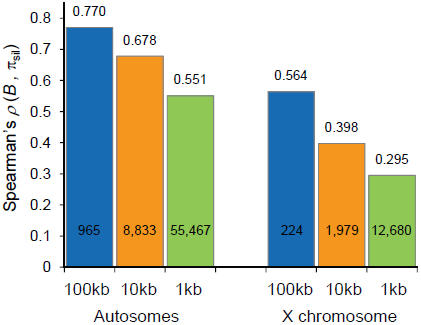
Correlation coefficients between estimates of *B* and levels of polymorphism at noncoding sites (π_sil_) for different physical scales. Spearman’s rank correlation coefficients (*ρ*) based on the analysis of 100-, 10- and 1-kb non-overlapping regions are shown above columns (*P* < 1×10^−12^ in all cases) and the number of regions analyzed is shown within columns. Data shown for the default model M_LN,StdMut,CO+GC_.

**Table 1.** Correlation coefficients between estimates of *B* and levels of polymorphism at noncoding sites (**π**_sil_) for different BGS models.

| | | DDFE | $U$ | Recombination | Spearman's rank correlation $\rho^a$ | |
| --- | --- | --- | --- | --- | --- | --- |
|  |  |  |  |  | Complete Chromosomes | Trimmed Chromosomes |
| **BGS Model** | $M_{\text{LN,StdMut,CO+GC}}$ | Log-normal | 1.2 | CO+GC | 0.770 (0.836) | 0.529 (0.655) |
| | $M_{\text{G,StdMut,CO+GC}}$ | Gamma | 1.2 | CO+GC | 0.772 (0.838) | 0.531 (0.659) |
| | $M_{\text{LN,LowMut,CO+GC}}$ | Log-normal | 0.6 | CO+GC | 0.770 (0.836) | 0.529 (0.655) |
| | $M_{\text{G,LowMut,CO+GC}}$ | Gamma | 0.6 | CO+GC | 0.773 (0.838) | 0.531 (0.659) |
| | $M_{\text{LN,StdMut,CO}}$ | Log-normal | 1.2 | CO | 0.749 (0.808) | 0.497 (0.598) |
| | $M_{\text{G,StdMut,CO}}$ | Gamma | 1.2 | CO | 0.761 (0.824) | 0.514 (0.630) |
| | $M_{\text{LN,LowMut,CO}}$ | Log-normal | 0.6 | CO | 0.749 (0.808) | 0.497 (0.596) |
| | $M_{\text{G,LowMut,CO}}$ | Gamma | 0.6 | CO | 0.761 (0.824) | 0.514 (0.630) |
| **Local Crossover** | $c$ | | | CO | 0.677 (0.732) | 0.397 (0.486) |

Spearman's rank correlation coefficient ( $\rho$ ) between estimates of $B$ and $\pi_{\text{sil}}$ ( $P < 1 \times 10^{-12}$ for all cases). Also shown, $\rho$ between local crossover rates ( $c$ ; cM/Mb) and $\pi_{\text{sil}}$ ( $P < 1 \times 10^{-12}$ ). Results shown based on the analysis of non-overlapping 100-kb autosomal regions.
<sup>a</sup> Values in parenthesis indicate the highest $\rho$ between $B$ and $\pi_{\text{sil}}$ along a single chromosome arm.

The predictive nature of the *B* landscape in *D. melanogaster* remains remarkably high along trimmed autosomes where BGS has been often assumed to play a minor role explaining variation in levels of polymorphism. The correlation between *B* and π_sil_ is *ρ* = 0.529, ranging up to *ρ* = 0.655 when trimmed chromosome arms are analyzed separately (*P* < 1×10^−12^ in all cases). Additionally, the BGS models investigated generate a stronger association between *B* and π_sil_ than between estimates of crossover (*c*) and π_sil_, particularly along trimmed chromosomes (*ρ* = 0.677 and *ρ* = 0.397 for complete and trimmed autosomes, respectively). This last result exposes the limitations of using local *c* as an estimate for the overall strength of linked selection at a given genomic position, and highlights the importance of including long-range information of recombination rates and gene structures. Altogether, these results show the high predictive value of simple BGS models, with almost 60% of the observed variance in π_sil_ across 100-kb autosomal regions explained by BGS, a percentage that is as high as ∼70% when investigating variation in nucleotide diversity along individual chromosome arms (see Table 1 and below for analyses at finer physical scale).

#### Robustness to different BGS parameters

The results presented above suggest that BGS could explain a very large fraction of the observed variation in diversity across the genome under the model that fits better our current selection and mutation data in *D. melanogaster* (model M_LN,StdMut_),. Importantly, BGS models using different DDFEs (M_G_ instead of M_LN_) and/or a lower deleterious mutation rate (M_LowMut_ instead of M_StdMut_) generate equivalent results. Table 1 shows *ρ* between *B* and π_sil_ for the different BGS models investigated: *ρ* between *B* and π_sil_ ranges between 0.749 and 0.773 for complete autosomes, and between 0.514 and 0.531 for trimmed autosomes (*P* < 1×10^−12^ in all cases). The study of intergenic and intronic sites separately generates similar outcomes, with *ρ* between *B* and π_intergenic_ ranging between 0.736 and 0.739, and *ρ* between *B* and π_intron_ ranging between 0.741 and 0.744 for the different models (*P* < 1×10^−12^ in all cases). The similarity of outcomes should not be surprising based on the very high pairwise rank correlations between estimates of *B* generated by the different BGS models described above (Table S3). Taken together, these results emphasize the robustness of the approach to generate genome-wide baselines of *B* to study variation in nucleotide diversity along chromosomes.

#### The X chromosome

Langley et al. (2012) [26] recently showed that the correlation between crossover rates and levels of polymorphism is weaker along the X chromosome than in autosomes. In agreement, we also observe that the association between *B* and π_sil_ along the X chromosome is weaker than in autosomes, although it is still highly significant. Estimates of *ρ* between *B* and π_sil_ range between 0.564 and 0.568 for the different BGS models (*P* < 1×10^−12^ in all cases), and between 0.366 (*P* = 4×10^−7^) and 0.373 (*P* = 2×10^−7^) for the trimmed X chromosome. Moreover, π_sil_ shows a weaker correlation with crossover rates than with *B* also along the X: *ρ* between π_sil_ and *c* is 0.526 (*P* < 1×10^−12^) and 0.322 (*P* = 9.1×10^−6^) for the complete and trimmed X chromosome, respectively. These results are consistent with BGS playing a weaker role along the X chromosome due to higher recombination rates (see above and [26,30,33,61]. As discussed below, however, additional causes might be also influencing X-linked extant variation, including higher effectiveness of selection relative to autosomes.

### Candidate regions for recent selective sweeps or balancing selection using BGS predictions as baseline

The robustness and high predictive power of the BGS models to explain qualitative trends of nucleotide diversity across the genome, suggest that we can investigate the presence of other forms of selection by searching for regions that depart from BGS expectations. We, therefore, compared observed π_sil_ and levels of diversity predicted by *B,* and parameterized departures by using studentized residuals (π_sil-R_; see **Material and Methods**). Overall, the distribution of π_sil-R_ does not show a significant departure from normality (χ^2^ = 28.9, d.f.= 23, *P* = 0.183) thus validating the approach. Nevertheless, there are 58 outlier regions with nominal *P* < 0.05, 24 regions with significantly negative π_sil-R_ (revealing a deficit in π_sil_ relative to BGS expectations) and 34 regions with significantly high π_sil-R_ (revealing a relative excess of π_sil_ ).

Regions with a relative deficit of π_sil_ are candidate regions for recent adaptive events [2,8,63] and our data confirms the presence of several regions with the fingerprints of a recent selective sweep across the *D. melanogaster* genome [22,23,25,26,29]. The strongest signal of selection at this 100-kb scale, and the only region that shows a departure that remains significant after correction for multiple tests [*P* < 1.6 × 10^−6^, with false discovery rate (FDR) *q*-value < 0.10], suggests a recent selective sweep at position 8.0-8.1 Mb along chromosome arm 2R. This genomic region includes gene *Cyp6g1* and also showed the strongest signal of directional selection and selective sweep in large-scale population genetic analyses of North American [26] and Australian *D. melanogaster* [64] populations. Not all regions with a severe reduction in π_sil_ across the trimmed genome, however, may need recent adaptive explanations. A number of regions with π_sil_ much smaller than the median (e.g., 0.002 relative to a median of 0.008; see Figure 6A) also show estimates of *B* of 0.25 or smaller and, thus, the observed π_sil_ would be close to the predicted level of neutral diversity when a BGS context is taken into account.

**Figure 6.**
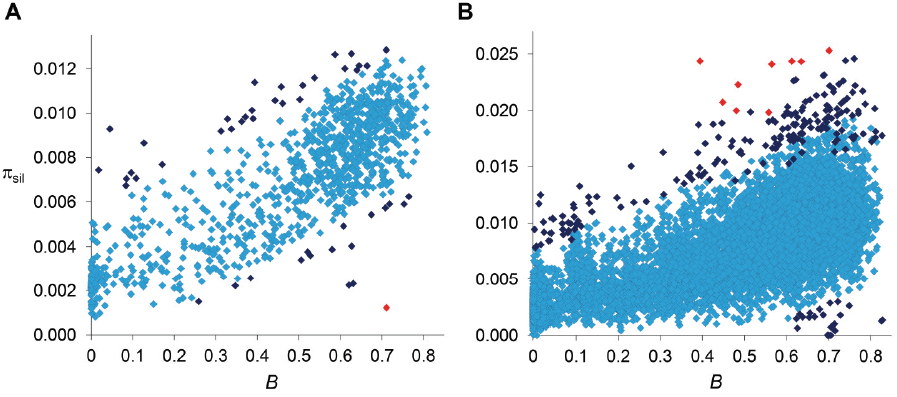
Relationship between estimates of *B* and π_sil_ for 100- and 10-kb regions. **(A)** Relationship between estimates of *B* and π_sil_ for 100-kb autosomal regions. Spearman’s *ρ* = 0.770 (1,189 regions, *P* < 1×10^−12^). **(B)** Relationship between estimates of *B* and π_sil_ for 10-kb autosomal regions. Spearman’s *ρ* = 0.678 (8,883 regions, *P* < 1×10^−12^). Dark blue and red diamonds indicate regions with π_sil_ significantly different (higher or lower) than predicted based on residual analysis: dark blue (*P* < 0.01), red (FDR-corrected *q* < 0.10). Data shown for the default model M_LN,StdMut,CO+GC_.

#### Uncovering the signature of balancing selection

Numerous regions show an excess of π_sil_ relative to the BGS baseline and are, therefore, candidates for containing sequences experiencing balancing selection. Importantly, many of the regions showing significantly higher π_sil-R_ would not stand out unless when compared to region-specific BGS predictions. At this scale, for instance, the second strongest departure from BGS expectations is located at position 18.4-18.5 Mb along chromosome arm 3L (*P* = 1×10^−4^), with a π_sil_ of 0.0093 that would not be noticeably high using the standard approach of comparing it to genome-wide expectations (Figure 6A). Another candidate region under balancing selection is located in cytological location 38A, showing a level of diversity also close to the average along autosomes but almost three-fold higher than the level predicted by *B (P* = 0.002). This candidate genomic region with a relative excess of π_sil_ is within a QTL detected for variation in life span and proposed to be maintained by balancing selection in *D. melanogaster* [65–67]. Certainly, the detection of many regions with a relative excess of π_sil_ based on BGS expectations opens the possibility that molecular signatures of balancing selection may have been masked by BGS effects when compared to purely neutral expectations without linkage effects, and thus be more prevalent than currently appreciated.

#### Analyses of BGS effects and outliers of diversity at 10-kb and 1-kb scales

Because the genome-wide recombination maps used in this study were generated to have good accuracy at the scale of 100-kb, our BGS models assumed homogeneous rates within each 100-kb region. Notably, these models predict variation in *B* across 100-kb regions due to the heterogeneous location of genes and exons within these regions and the differential effects of proximal and distal flanking regions. Nevertheless, detailed analyses of a few small genomic regions have revealed recombination rate variation at a smaller scale [32,68]. Therefore, outliers from BGS expectations at scales smaller than 100-kb can reveal the localized fingerprints of other types of selection (directional or balancing selection) but the possibility of uncharacterized heterogeneity in recombination within these regions cannot be formally ruled out. That said, the study of the relevant size of the genomic region adding up BGS effects at a given focal point (with D*_B75_* > 200 kb; see above), suggests that very local recombination variation may play a limited role influencing *B* at a focal point. With these caveats and considerations in mind, we investigated the presence of outliers at the scale of 10 and 1-kb to identify *candidate* regions under positive or balancing selection using the approach discussed for 100-kb regions.

The strong relationship between *B* and the observed level of silent diversity is maintained when analyzing smaller regions (Figure 5). At 10-kb scale, *B* remains a very good predictor of π_sil_ along complete autosomes (*ρ* = 0.678, 8,883 regions; Figure 6B) whereas *ρ* is 0.551 (55,467 regions) at the finest scale of 1-kb (*P* < 1×10^−12^ in both cases). The use of BGS models with different parameters (M_G,StdMut_, M_G,LowMut_ or M_LN,LowMut_) generate virtually equivalent results, with *ρ* between estimates of *B* and π_sil_ ranging between 0.678 and 0.682 for analyses at the 10-kb scale, and with *ρ* ranging between and 0.551 and 0.554 for analyses at the 1-kb scale. As observed before, *B* along the X chromosome shows reduced association with π_sil_ than for autosomes also at 10- and 1-kb resolution. For X-linked regions, the correlation between *B* and π_sil_ is *ρ* = 0.397 (1,979 regions) and 0.295 (12,680 regions) for 10- and 1-kb regions, respectively (*P* < 1×10^−12^ in both cases).

*Outliers at 10-kb scale*: Among the 10,812 regions under analysis, 208 depart from expectations with nominal *P* < 0.01, 20 showing a relative deficit of π_sil_ and 188 with a relative excess of π_sil_. Nine of these regions remain significant after correction for multiple tests (FDR *q*-value < 0.10), and all of them show an excess of π_sil_ (Figure 6B). Among the candidate regions for a recent selective sweep (relative deficit of π_sil_), we detect 3 genomic regions with clusters of several 10-kb regions with nominal *P* < 0.01. In agreement with the analyses at 100-kb scale, 10 consecutive 10-kb regions show significant deficit of variation, near gene *Cyp6g1* (see above and [26,64]). A second cluster of six 10-kb regions with reduced variation within a 120-kb interval is detected in chromosome arm 2R, with a peak signal centered at gene *Dll* (*Distalless*), a transcription factor that plays a role in larval and adult appendage development. A third region with two adjacent 10-kb regions showing a relative deficit of π_sil_ is located in the X chromosome, centered at gene CG32783 (a protein-encoding gene with no known orthologs outside the *melanogaster* subgroup).

On the other hand, we detect many 10-kb regions that show a relative excess of diversity and are candidate regions for balancing selection. The strongest signal associated with excess of π_sil_ based on 100-kb regions now shows significantly higher variation at four adjacent 10-kb regions (19.360-19.400 Mb along chromosome arm 2L). Other strong candidate regions under balancing selection detected at this fine scale (all with FDR *q*-value < 0.10) include the genes PH4αNE2 (oxidation-reduction process), *dpr20* (sensory perception of chemical stimulus), *Dhc16F* (regulation of transcription and microtubule-based movement) and *Gαi*. Note that gene *PH4αNE2* was also shown to have an excess of protein polymorphism in the sister species *D. simulans* [24], thus potentially revealing a rare case of trans-specific balancing selection.

To further investigate whether the outlier regions are indeed associated with recent selective sweeps and balancing selection, we estimated Tajima’s *D* (*D*_T_; [69]) to quantify potential differences in the frequency of polymorphic variants within the population. Selective sweeps and balancing selection are predicted to generate an excess of low-frequency (negative *D*_T_) and intermediate-frequency (positive *D*_T_) variants, respectively. In agreement, outlier regions with negative and positive π_sil-R_ have more negative (Mann-Whitney *U* Test, *P* = 4.3×10^−10^) and more positive (*U* test, *P* < 1×10^−12^) *D*_T_, respectively, than regions not departing from BGS expectations. Moreover, we observe a positive association between π_sil-R_ and *D*_T_ across the genome (*ρ* = 0.280, *P* < 1×10^−12^) that is stronger than between π_sil_ and *D*_T_ (*ρ* = 0.087). Notably, this associating between π_sil-R_ and *D*_T_ increases across trimmed chromosomes (*ρ* = 0.302 and *ρ* = 0.637 for trimmed autosomes and the X chromosome; *P* < 1×10^−12^). Taken together, these results reinforce the concept that estimates of π_sil-R_ obtained when using BGS predictions as baseline are a good predictor of recent selective sweeps and balancing selection, capturing departures in number of polymorphic sites as well as the expected consequences on variant frequency.

*Outliers at 1-kb scale*: The study of 1-kb regions reveals 1,213 of them having diversity levels that depart from BGS expectations with nominal *P* < 0.01, 1,186 and 27 of these regions show a relative excess and deficit of π_sil_, respectively. Fifty-two regions (0.076%) show an FDR corrected *q* < 0.10, all with higher π_sil_ than predicted by the BGS model. Twenty-five out of the 27 1-kb regions showing a reduction in π_sil_ with *P* < 0.01 cluster together at position 8.017 − 8.103 Mb along chromosome arm 2R, in agreement with the analyses at 100- and 10-kb scales that detected outlier regions near gene *Cyp6g1*. This more detailed analysis suggests that the target of selection is likely located at or proximal to *Cyp6g1*. The other two outlier regions with deficit in π_sil_ are located at genes *cic* (regulation of transcription) and CG11266 (mRNA binding and splicing).

In terms of regions with an excess of π_sil_, the strongest signal of balancing selection based on 100- and 10-kb analyses (19.3-19.4 along chromosome arm 2L) is now localized with more precision close to genes CG17349 and CG17350. Other regions detected at 10-kb scale are also identified at this finest scale and include genes *PH4*α*NE2*, *dpr20*, and *Dhc16F* (all with FDR *q*-value < 0.10). Additional candidate regions under balancing selection include genes *CecA1*/*CecA2*/*CecB* (antibacterial humoral response), *Sema*-5c (olfactory behavior), CG5946 (inter-male aggressive behavior), *IM4* (defense response), *Cyp6a16* (a P450 gene), and three genes encoding cuticular proteins (*Cpr11A*, *Cpr62Bb* and *Cpr65Ec*), among others. Finally, the study of 1-kb regions also shows a strong positive relationship between π_sil-R_ and *D*_T_ (ρ = 0.477, *P* <1×10^−12^), in agreement with the expected consequences of selective sweeps and balancing selection on the frequency of segregating variants (see above).

### Negative relationship between estimates of *B* and the rate of protein evolution.

Another prediction of the models of selection and linkage (either HHss or BGS) is that, parallel to a reduction in intra-specific variation, there will be a reduction in efficacy of selection (i.e., the Hill-Robertson effect [10,70–76]). This general prediction has been previously confirmed in *Drosophila* using local low resolution crossover rates as indirect measure for the magnitude of Hill-Robertson effects acting on a gene. These studies showed weak but highly significant associations between crossover rates and estimates of codon usage bias or rates of protein evolution [71,74,75,77–85].

To investigate whether *B* landscapes also capture differences in efficacy of selection, we focused on selection against amino acid substitutions along the *D. melanogaster* lineage, after split from the *D. simulans* lineage less than 5 mya [86]. To this end, we obtained the ratio of nonsynonymous to synonymous changes (*ω*, *ω* *= d_N_/d_S_*) for 6,677 protein encoding genes and, more informatively, the variation in *ω* after controlling for selection on synonymous mutations based on residual analysis (*ω*_R_; see **Materials and Methods** for details). When each gene is analyzed as a single data point, there is a negative association between *B* and *ω*_R_ (*ρ* = − 0.086, *P* = 2×10^−12^; Table 2). Interestingly, and contrary to the results of nucleotide diversity, the X chromosome shows a tendency for a stronger effect of *B* on *ω*_R_ than autosomes: *ρ* = − 0.189 (*P* = 3.4×10^−8^) and *ρ* = − 0.071 (*P* = 5.7×10^−8^) for X-linked and autosomal genes, respectively. An equivalent but more clear pattern is observed at the scale of the resolution of our recombination maps (100 kb), where estimating the average *ω* and *ω*_R_ for all genes within each region also allows for reducing idiosyncrasies of different genes influencing rates of protein evolution (e.g., gene expression breadth and levels, protein length, etc.; see [84]). At this scale, variation in *B* is negatively associated with *ω*_R_ along autosomes (*ρ* = −0.160, *P* = 6.1×10^−6^) and the X chromosome (*ρ* = − 0.367, *P* = 1.5×10^−6^; Table 2). Again, the association between estimates of *B* and rates of protein evolution is robust to different BGS models and parameters. Equivalent *ρ* are obtained for all eight BGS models investigated, and this is observed when analyzing individual genes (*ρ* between *B* and *ω*_R_ ranging between − 0.0856 and − 0.0874; *P* ≤ 3×10^−12^) and when using the average *ω*_R_ for genes within 100-kb regions (*ρ* ranging between − 0.187 and − 0.193; *P* ≤ 5.9×10^−9^) across the whole genome.

**Table 2.** Correlation coefficients between estimates of *B* and rates of protein evolution (****ω****_R_).

|  |  | **Spearman's $\rho$** | **Probability** |
| --- | --- | --- | --- |
| **Individual genes** | All | - 0.057 | $2 \times 10^{-12}$ |
| | Autosomes | - 0.071 | $5.7 \times 10^{-8}$ |
| | X chromosome | - 0.189 | $3.4 \times 10^{-8}$ |
| **100-kb regions** | All | - 0.187 | $6.6 \times 10^{-9}$ |
| | Autosomes | - 0.160 | $6.1 \times 10^{-6}$ |
| | X chromosome | - 0.367 | $1.5 \times 10^{-6}$ |

Estimates of *B* were obtained from model M_LN,StdMut,CO+GC_ whereas estimates of the rate of protein evolution for each protein encoding gene (*ω*_R_) were obtained after controlling for selection on synonymous mutations based on residual analysis (see **Materials and Methods** for details). Spearman’s rank correlation coefficients (*ρ*) are shown between B and *ω*_R_ for individual genes and for the average *B* and *ω*_R_ for all genes within 100-kb non-overlapping regions.

### Temporal variation in recombination landscapes and its consequences

The association between *B* and rates of amino acid substitution along the *D. melanogaster* lineage, although highly significant in terms of associated probability, is much weaker than that for levels of polymorphism at silent sites. Heterogeneity in overall evolutionary constraints among proteins is expected to add substantial variance when investigating the consequences of *B* on *ω* relative to studies of *B* on π_sil_ because π_sil_ is only influenced by local *N_e_* and the mutation rate. Nevertheless, temporally variable recombination rates at a given genomic location would also reduce the association between *B* and *ω*. Indeed, the high degree of intra-specific variation in crossover genetic maps within current *D. melanogaster* populations [31] together with differences in genetic maps between closely related *Drosophila* species [87–89] support the notion that recombination landscapes vary within short evolutionary scales, at least across trimmed chromosomes. Under this scenario, extant recombination rates and the corresponding estimates of linkage effects would be poor predictors of interspecific rates of protein evolution, even between closely related species.

In this study across the *D. melanogaster* genome, the use of recombination rates obtained experimentally would provide only an approximation for the relevant *B* along the lineage leading to *D. melanogaster* populations [84]. These estimates of *B* would be an even weaker predictor of *ω*_R_ (or *ω*) along the *D. simulans* lineage after split from the *D. melanogaster* lineage. In agreement, we observe no significant relationship between *B* and *ω*_R_ estimated along the *D. simulans* lineage (*ρ* = −0.009 based on the default BGS model whereas the other BGS models generate *ρ* ranging between -0.014 and +0.011; *P* > 0.25 in all cases). A similar result has been obtained in comparisons of crossover rates and rates of protein evolution between two other closely related *Drosophila* species, *D. pseudoobscura* and *D. persimilis* [88].

In species where BGS plays a significant role, temporal fluctuations in recombination landscapes could influence a number of analyses of selection that assume constancy in *N_e,_* including estimates of the fraction of adaptive substitutions (α; [6,90–92]). Several studies have shown that the bias in estimating α can rapidly reach considerable values as a consequence of demographic changes, with α being overestimated when *N_e_* influencing polymorphism is larger than *N_e_* influencing divergence (*N_e_Pol_* > *N_e_Div_*; [34,91]). We propose that temporal changes in recombination at a given genomic position would generate an equivalent scenario, with a *B* influencing polymorphism (*B__Pol_*) that differs from long-term *B* influencing divergence (*B__Div_*), or the corresponding terms for local *N_e_*. Because long term *N_e_* can be approximated by its harmonic mean [93], temporal fluctuations of recombination landscapes (and of local *B* and, therefore, local *N_e_*) would also predict a tendency for local *N_e_Pol_* ≥ *N_e_Div_*. Such scenario would allow amino acid changes to make a larger relative contribution to divergence than to polymorphism, particularly in regions where recombination has recently increased, and thus bias estimates of α upward.

Precise quantitative predictions of the potential bias in α would minimally depend on the rate, magnitude and physical scale of the variation in recombination landscapes along lineages. To obtain initial insight into the potential effects of temporal changes in recombination rates on estimates of α, we investigated a rather simple and conservative scenario with forward population genetic simulations. In particular, we used the program SLIM [94] to capture the consequences of temporal changes in linkage effects on estimates of α when only neutral and deleterious mutations occur along an archetypal 1-Mb region for *D. melanogaster* that includes 100 protein coding genes (see **Materials and Methods** for details). Figure 7 shows the results of estimating α at selected sites under fluctuating recombination rates, with cycles of moderately high recombination for 1*N* generations (with *N* indicating the diploid population size) followed by moderately low (not zero) recombination for 3*N* generations. Estimates of α based on models that assume constant population size [34,91,95] overestimate the true α particularly when extant recombination is high (α > 0.3), with an overall α ∼ 0.15 when the data is combined from all temporal points. As expected, models allowing for population size change [91] generate more unbiased estimates of α that show, nonetheless, a tendency upward, likely due to limitations assessing older population size changes.

**Figure 7.**
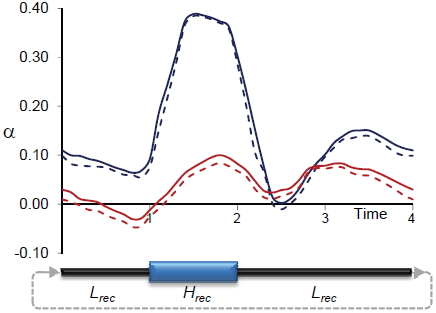
Estimates of α due to temporally fluctuating recombination rates and variable BGS effects. Results based on forward population genetic simulations of 10,000 diploid individuals (*N*), a chromosome segment of 1 Mb containing 100 genes, and two types of mutations: neutral and deleterious (see **Materials and Methods** for details). Cycles of fluctuation recombination followed phases of moderately high recombination for 1*N* generations (*H*_rec_ phase) and low recombination for 3*N* generations (*L*_rec_ phase). Estimates of α at selected sites obtained following the models proposed by Eyre-Walker and Keightley [91,95] every 0.1*N* generations. Blue and red lines indicate estimates of α assuming constant population size and variable population sizes, respectively. Continuous and dashed lines indicate estimates of α with and without correction for the effect of polymorphism to divergence, respectively.

## DISCUSSION

Discerning the relative contribution of different types of selection to patterns of intra- and interspecific variation is not trivial, in part because essential population and demographic parameters are often not known and the potential interactions among them are still poorly characterized. Here, we obtained the baseline of diversity across the genome predicted by BGS models completely independent of the data on nucleotide variation. The results of this study suggest that there might be no euchromatic region completely free of linkage effects to deleterious mutations in *D. melanogaster*. Instead, there are only genomic sites (neutral or under selection) associated with diverse degrees of BGS effects and thus under highly variable local *N_e_* even across the recombining regions of the genome. In this regard, the heterogeneity in local *N_e_* may bias population genetic estimates of selection or recombination rates if such variation in not taken into account. The pervasive presence of BGS has also potential consequences on demographic studies because BGS is known to generate a moderate excess of low-frequency variants [96–102]. As a result, demographic inferences based on allele-frequencies may be skewed, with consistent patterns suggestive of a recent population expansion.

The next question investigated was how much of the patterns of variation in *D. melanogaster* could be explained by purifying selection alone. The results are consistent with BGS playing a major role in explaining the observed heterogeneity in nucleotide diversity across the entire *D. melanogaster* genome. At 100-kb scale, BGS can explain ∼58% (*ρ* = 0.749 − 0.773 for different BGS models) of the variation of π_sil_ across the genome. The study of smaller regions reduces the statistical association between *B* and π_sil,_ but even when analyzing 10- and 1-kb regions, *B* explains ∼46% (*ρ* = 0.678 − 0.682), and ∼30% (*ρ* = 0.551 − 0.554) of all the observed variance in π_sil_, respectively. These percentages increase up to ∼70% (*ρ* = 0.836) for 100-kb regions, ∼53% (*ρ* = 0.728) for 10-kb regions, and ∼36% (*ρ* = 0.599) for 1-kb regions, across individual chromosome arms.

Importantly, these conclusions are robust to differences in parameters of the BGS model (e.g., deleterious mutation rates, DDFE, or recombination) even though the precise magnitude of BGS effects does depend on these parameters. Median estimates of *B* range between 0.337 and 0.769 for different BGS models, but the distribution of *B* along the chromosomes maintains the same rank order (pairwise *ρ* between estimates of *B* generated by the BGS models is ≥ 0.9856) and is similarly associated with the observed distribution of nucleotide diversity.

Taken together, the results and conclusions of this study imply that null expectations based on models that ignore linkage effects to deleterious mutations (or assume homogeneous distribution of *N_e_* across the genome) are likely inaccurate in *D. melanogaster,* even across trimmed chromosomes. The fact that deleterious mutations are more frequent than mutations involved in balancing selection or adaptive events, together with the robustness of our conclusions to reasonable ranges of selection and mutation parameters, suggest that BGS predictions may be adequate as a baseline of diversity levels and can be used detect outlier regions subject to other selective regimes.

The use of *B* as baseline reveals several regions with signal of recent directional selection and associated selective sweeps, where levels of variation are lower than those predicted by BGS alone (e.g., near gene *Cyp6g1* [26,64]). In agreement with expectations after a selective sweep, these regions also show an excess of variants at low frequency. A number of other regions with reduced levels of variation, however, are located in recombining genomic regions where BGS is predicted to have strong effects. These results, therefore, are consistent with a detectable but limited incidence of recent classic ‘hard sweeps’ (where diversity is fully removed near the selected sites) caused by beneficial mutations with large effects. Still it must be acknowledged that older selective sweeps, sweeps caused by weakly beneficial mutations, or ‘soft sweeps’ associated with standing genetic variation or polygenic adaptation [103,104] would not be detected in our study.

In fact, estimates of the proportion of adaptive substitutions, α, indicate that beneficial mutations are not rare in *Drosophila* [6,90–92]. Moreover analyses of nucleotide diversity around amino acid substitutions suggest that a majority of these beneficial mutations involve small effects on fitness [25,105,106] and cause detectable but very localized reduction in adjacent diversity (at the scale of 25 bp; [105,106]). Therefore, a bulk of beneficial mutations in *Drosophila* may be difficult to detect in genome-scans of variation but could contribute significantly to differences between species and overall adaptive rates of evolution.

BGS baselines, however, are particularly adequate to detect genomic regions under balancing selection because the predicted genomic signature of an excess of diversity may not be always evident when compared to genome-wide averages or purely neutral expectations (see Figure 6). Indeed, the use of a *B* baseline across the whole *D. melanogaster* genome provides the adequate local *N_e_* context caused by purifying selection and corresponding linkage effects. In all, our study uncovers numerous candidate regions for balancing selection, identifying genes involved in sensory perception of chemical stimulus, antibacterial humoral response, olfactory behavior, inter-male aggressive behavior, defense response, etc. These genomic regions not only show a relative excess of polymorphic sites but also have segregating variants at higher frequency than the rest of the genome (another telltale sign of balancing selection). The results based on the study of a single population are appealing since heterogeneous environments or temporal changes predict the maintenance of local polymorphism through balancing selection [107–109], and are consistent with analyses of clines in *D. melanogaster* that detected the signature of local adaptation and spatially varying selection between populations [64,110,111].

Additionally, the results evidence a clear difference between autosomes and the X chromosome in terms of the consequences of variable *B* on levels of variation. While the X chromosome exhibits a reduced association between *B* and neutral diversity than autosomes, it shows a better fit between *B* and rates of protein evolution and efficacy of selection. These patterns are predicted by higher rates of recombination and a higher fraction of deleterious mutations with minimal role generating BGS effects in the X (see **Material and Methods**). Another factor possibly influencing this difference between X and autosomes is stronger efficacy of selection in the X chromosome [112]. Indeed, events of adaptive and/or stabilizing selection would distort levels of neutral diversity at linked sites beyond BGS predictions and stronger selection in the X would explain a reduced association between *B* and levels of diversity along the X relative to autosomes. Such combined scenario of reduced BGS effects and stronger selection would also explain a number of patterns observed in *Drosophila*, including an increased degree of synonymous codon usage bias [112–115], stronger purifying and positive selection acting at the level of protein evolution in X-linked genes [115,116], and the ‘faster-X’ effect [117,118] reported at both protein and gene expression levels [119–123]. Of interest will be the study of species with higher average recombination rate in autosomes than in the X, thus partially uncoupling differences in linkage effects and X-specific patterns.

Moreover, the results of this and previous studies [26,87–89] suggest that recombination rates change frequently in the *Drosophila* genus, and often involve differences in the distribution of recombination rates across the genome (i.e., a change of the recombination landscape). Very recent changes in recombination landscapes would uncouple extant recombination rates and polymorphism patterns generated during the last few ∼*N_e_* generations and, therefore, the reported contribution of BGS to patterns of diversity across the genome may be underestimated. On the other hand, changes in the recombination environment in species where BGS plays a significant role would also predict temporal variation in local *B* and impact a number of studies of selection that assume constancy of *N_e_* along lineages or that changes in *N_e_* should be equivalent across the genome.

Temporally variable recombination landscapes can generate, for instance, spurious evidence for multimodality in the distribution of fitness effects, or lineage-specific physical clustering of amino acid changes (such clustering has been observed among *Drosophila* species [124,125]). Another consequence of temporally variable recombination landscapes (and local *B* and *N_e_*) would be gene- or region-specific inequality of short- and long-term *N_e_*. Regions that have recently increased in recombination would show patterns of variation suggesting population expansion or bias estimates of α upward, making it less negative or even positive with no adaptive mutations [126]. Moreover, these changes in recombination landscape would also forecast substantial between-gene variation in α without requiring adaptive evolution. At this point, therefore, there is the open possibility that positive estimates of α in *Drosophila* and other species with large population size (see [6,22] and references therein) may be influenced, to an unknown degree, by temporal changes in recombination rates and landscapes. Future analyses designed to estimate population size changes or the strength and frequency of adaptive events would, therefore, benefit from including variable BGS effects across genomes and along lineages, ideally discerning local variation in BGS (and local *N_e_*) from genome-wide patterns that may represent true demographic events. Genome-wide analyses may also need to consider the non-negligible presence of regions under balancing selection to prevent overestimating the extent of recent sweeps based on a relative reduction in levels of diversity.

Finally, a number of limitations and areas for future improvement need to be mentioned. In terms of selection, we investigated BGS models based on either a log-normal or a gamma DDFE [39–43,46,127]. But trying to include all possible mutations with deleterious effects across a genome into a single distribution is a clear oversimplification. A combination of different DDFEs for different groups of genes and/or sites would be a more fitting approach, ideally based on *a priori* information independent of levels of variation (e.g., based on amino acid composition, expression levels and patterns, connectivity, etc. [84]).

To gain initial insight into the potential consequences of more realistic models, we obtained estimates of *B* under BGS models that allow for two DDFEs (one for nonsynonymous changes and one for changes at constrained noncoding sites; see **Materials and Methods** for details). We then investigated whether such models would alter our conclusions, mostly in terms of the proposed adequacy of using BGS predictions as baseline to detect outliers. The results show that models with two DDFEs depart only marginally from the models with a single DDFE: rank correlations between estimates of *B* predicted by these hybrid models and all eight previous models show *ρ* ranging between 0.946 and 0.998. The association between predicted *B* and the variation of π_sil_ across the genome is also very high (albeit slightly lower than for models based on a single DDFE), with *ρ* ≥ 0.711 at 100-kb scale and *ρ* ≥ 0.504 at 1-kb scale. Notably, the comparison of outliers generated by these 2-DDFE models with those obtained by the models described above reveals few differences. For instance, the study of a model assuming a log-normal DDFE for nonsynonymous mutations and a gamma for deleterious noncoding mutations (M_LN/G,StdMut,CO+GC_) shows that 50 out of the 52 1-kb outlier regions after correcting for multiple testing are also predicted to be equally significant outliers based on this hybrid and most different model (the other two regions show departure with *P* < 0.0001). Overall, 87.1% of the 1,213 1-kb regions showing departure at *P* < 0.01 using the default BGS model are also detected as outliers (*P* < 0.01) under this hybrid model. These results further support the robustness of the proposed approach to detect outlier regions based on BGS baselines.

Another line of future improvement is the physical resolution of our recombination maps and, perhaps more important, how well these maps represent the recent history of a population or species. To generate *B* landscapes, we used recombination maps that were experimentally obtained (independent of nucleotide variation data) and capture some intra-specific variation in recombination landscapes (see [31] for details). The fact that these *B* landscapes explain a very large fraction of the variance in nucleotide diversity across the genome suggests that these recombination maps represent quite accurately the recombination landscape in the recent history of *D. melanogaster*. Future recombination maps obtained from additional natural strains and populations, or under different biotic/abiotic conditions, can only improve the confidence in outlier regions and, ultimately, our understanding of selective and demographic events in this species.

## MATERIALS AND METHODS

### General BGS expectations

The consequences of BGS at a given nucleotide position in the genome (focal point) can be described by *B* [12–15], the predicted level of neutral nucleotide diversity under circumstances where selection and linkage are allowed (π) relative to the level of diversity under complete neutrality and free recombination (π**_0_**). Following [15,16], we have

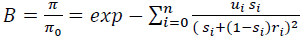

where *u_i_* is the deleterious mutation rate at the *i-*th selected site out of the *n* possibly linked sites, *s_i_* indicates the selection coefficient against a homozygous mutation and *r_i_* is the recombination frequency between the focal neutral site and the selected *i-*th site. Note that under a BGS scenario *B* can only be equal (no BGS effects) or lower than 1, and *B* << 1 indicates strong BGS effects.

Molecular evolutionary analyses indicate variable fitness effects of deleterious mutations [44,121,127,128] and, therefore, a probability distribution of deleterious fitness effects (DDFE) of mutations at site *i*, *ϕ(s_i_*) needs to be included, generating

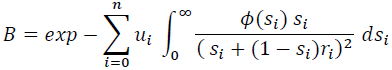

*B,* therefore, is predicted to vary across the genome as consequences of the known difference in recombination rates along chromosomes as well as due to the heterogeneous distribution of sites under selection across genomes (genes, exons, etc.). However, and because we are allowing the selection coefficient *s* to vary according to a distribution *ϕ(s*), some sites under selection may have selection coefficients too small to play any BGS effect, thus not all sites under section need to be included in the study [33]. We truncate the distribution of selection coefficients at *s*_T_ (*s*_T_ ∼ 1/ *N_e_*) because mutations with *s* < *s*_T_ are effectively neutral and do not contribute to BGS. Different DDFEs, therefore, will generate a different fraction of deleterious mutations with *s* > *s*_T_ and, therefore, different BGS effects (see below).

### High-resolution estimates of *B* across the whole *D. melanogaster* genome

We investigated a model that assumes the possibility of strongly deleterious mutations at nonsynonymous sites as well as at a fraction of noncoding sites, either untranslated genic (introns and 5’- and 3’-flanking UTRs) or intergenic regions. *B*, therefore, can be estimated at any focal neutral site of the genome as the cumulative effects of deleterious mutations at any other nonsynonymous, untranslated genic and intergenic site along the same chromosome. For simplicity’ and based on analyses of rates of evolution between *Drosophila* species [22,37,38,44,91], we assumed that the distribution and magnitude of selection on intergenic or UTR noncoding strongly selected sites [**ϕ**_*nc_ss*_(*s*)] is equivalent to that on nonysynonymous sites [**ϕ**_*aa*_(*s*)].

We can estimate *B* at any focal neutral site of the genome using the equation described above, **ϕ**_*aa*_(*s*), the actual genome annotation (*D. melanogaster* annotation release 5.47, http://flybase.org/) and high-definition recombination maps for crossing over and gene conversion events [31]. The analysis of every nucleotide sites along a single chromosomal arm taking into account all other sites under selection of this same chromosome would, however, require >1×10^13^ pair-wise nucleotide comparisons, each one requiring a numerical integration. To speed up the process we followed [17] and first obtained the integral

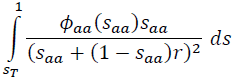

along a continuum for possible recombination rates between two sites, in our case for *r* ranging between 1×10^−10^ (equivalent to *c* = 0.01) to 1; we assumed *r* = 0 whenever the recombination distance between two nucleotides is smaller than 1×10^−10^. Because he recombination distance between any pair of nucleotides can be obtained, the generation of complete maps of *B* is now more tractable although still computationally very intensive. We then made the simplifying assumption of ignoring variation within 1-kb windows when estimating *B* at any another 1-kb window along the chromosome. That is, for the purpose of generating BGS effects, all sites within a 1-kb window have an equivalent recombination rate with sites within any other 1-kb window along the chromosome. This is a reasonable approximation at this time due to the fact that the resolution of our best genome-wide recombination maps is 100-kb and differences in recombination due to a few nucleotides play a minor role when comparing sites separated by tens of hundreds of kbs.

By dividing a chromosome arm into *L* adjacent 1-kb windows, *B* at a focal neutral site located at the center of the *j*-th window(*B_j_*), can be estimated by using:

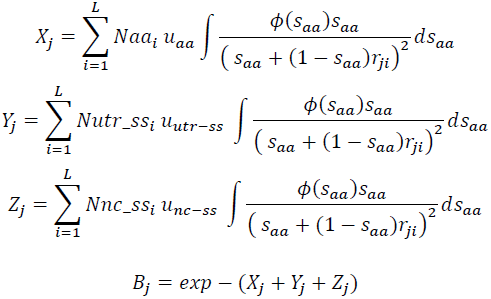

where *Naa_i_*, *Nutr_ss_i_* and *Nnc_ss_i_* are the number of nonsynonymous, UTR and intergenic sites possibly under strong selection within window *i*, respectively, and *r_ji_* is the recombination between the center of the focal window *j* and the center of window *i*, with *r_ji_* increasing in 1-kb intervals. *u*_*aa*_ , *u*_*utr-ss*_, and *u*_*nc-ss*_ are the deleterious mutation rate at nonsynonymous, UTR and intergenic sites , respectively. Note that this approach avoids the need to interpolate *B* estimates and generates a very detailed *B* landscape because all sites in the genome are actually being taken into account.

#### Deleterious mutation rates

Initial estimates of the mutation rate based on mutation accumulation lines in *D. melanogaster* suggested a mutation rate (*u*) of ∼ 8.4 × 10^−9^ /bp /generation, for point mutations and small indels [47]. More recent analyses suggest *u* ∼4-5 × 10^−9^ /bp/generation after removing data from a line with unusually high mutation rates [48–50]. Based on the fraction of conserved sites in exons and noncoding sequences [22,37], this lower mutation rate predicts a diploid rate of deleterious mutations per generation (*U*) of 0.6 for euchromatic regions. *U* ∼ 0.6, however, is an underestimate of the true *U* in *D. melanogaster* because it does not take into account the recent bottleneck experienced by North America populations (see [48]) or the deleterious consequences of TE insertions within genes and regulatory sequences. Additionally, the elevated mutation rate observed in one line (line 33 [47,48]) is not a new trait developed during the mutation accumulation experiment and, therefore, such genotype was present in the initial natural population of *D. melanogaster*. We, thus, use *U* = 0.6 as a lower boundary for the true deleterious mutation rate across the euchromatic genome (in models M_LowMut_).

TEs are pervasive across the *D. melanogaster* genome [51–60]. Cridland et al (2013) recently reported a detailed analysis of TE abundance and distribution in *D. melanogaster* based on deep-coverage, whole-genome sequencing [60]. Their analysis of the Drosophila Synthetic Population Resource (DSPR; [129]) showed a total of 7,104 TE insertions, with 633 TEs inserted within exons. To include the deleterious consequences of TE insertion, therefore, we obtained an approximate mutation (insertion) rate (*u*_TE_) based on mutation-selection balance predictions and the presence of TEs in genomic regions where they are most likely to cause deleterious effects (i.e., within exons). [We use TE presence in the DSPR lines instead of analyses based on the Drosophila Genetic Reference Panel (DGRP) due to the higher sequencing coverage in the DSPR lines, which increases the likelihood to detect and map TE insertions (see [60] for details)]. For non-recessive strongly deleterious mutations, mutation-selection balance predicts a probability of segregation *P_seg_* that is the product of the equilibrium frequency (*q*) and the number of chromosomes analyzed (*n*). *P_seg_* in exons (/bp) is 0.00003 (633 TEs / 21,000,000 exonic bp) and *q* = *P_seg_* / *n* = 2×10^−6^. Based on the very low frequency of segregating TEs in natural populations (i.e., most TEs in this study were present in a single genome in agreement with previous analyses of *D. melanogaster* populations; see [53,57,58]), we assume non-recessive deleterious effects and a heterozygous selection coefficient (*s_h_*) against TE insertions within exons to be equal or greater than for amino acid mutations (*s* ≥ 0.0025), so *u*_TE_ can be estimated by *u*_TE_ = *q s_h_,* with *u*_TE_ ≥ 5×10^−9^ /bp/generation. Thus, we obtain *U*_TE_ ∼0.6, a result in agreement with previous analyses of TE transposition rate that suggested that the genomic rate of TE transposition is of the same order as the nucleotide spontaneous mutation rate [55], and suggests *U* ∼1.2 when including point mutations, small indels and TE insertions across the euchromatic regions of the *D. melanogaster* genome.

#### Distribution of deleterious effects

The true probability distribution of selection coefficients against newly arising mutations is not known. Sensibly, no single deleterious distribution of fitness effects (DDFE) is likely to capture mutations at different genes or sites within genes. Different spatial and temporal conditions will, likely, further alter such DDFEs. As an approximation, we studied two DDFEs and the corresponding parameters proposed for *Drosophila* to partially capture the potential effects of different DDFEs on our results and conclusions: a gamma distribution [39–44] and a log-normal distribution [45,46], with the latter allowing to capture the presence of lethal mutations and fitting better *D. melanogaster* polymorphism data [45,46].

Following Loewe and Charlesworth [45], the log-normal DDFE has a median *s* of 0.000231 and standard deviation 5.308 for autosomes. A gamma DDFE for *Drosophila* is described by a shape parameter *k* = 0.3 and mean *s* of 0.0025 [33,38]. In both cases, we assumed a dominance coefficient *h* of 0.5. For models that include the contribution of TE insertions (*U* = 1.2), we further assumed that the DDFE of TEs is the same than of point mutations, and that the genomic sites where a TE insertion has deleterious effects are the same where point mutations and small indels have deleterious effects. Note that under the models that include a log-normal DDFE, the median *s* against individual TE insertions (0.000231) is equivalent to previous estimates in *D. melanogaster* of 0.0002 [54].

An interesting attribute of the log-normal distribution is that it predicts a higher frequency of extreme values, including effectively neutral (*s* < *s*_T_) and lethal (*s* > 1) mutations, than those predicted by a gamma distribution. Because effectively neutral and strongly deleterious mutations play a minimal role removing linked variation [130], a log-normal DDFE is predicted to generate weaker BGS effects than a gamma DDFE with similar median *s*. Finally, it is worth noting that the parameters for these two DDFEs were obtained independently of a BGS model and therefore are not expected to maximize BGS as explanation of the observed patterns of diversity across the genome while, at the same time, may represent underestimates of the true magnitude of selection.

#### BGS with crossover and gene conversion along D. melanogaster chromosomes

To estimate the recombination frequency between a focal neutral site and selected sites in BGS formulae, we used the whole-genome high-resolution recombination maps from *D. melanogaster* described in [31]. These maps describe crossing over and gene conversion rates along chromosome arms (except the small fourth chromosome) at 100-kb resolution. Following Langley et al ([131]; see also [45,132]), the total recombination frequency between a focal neutral site and a selected *i-*th site taking into account crossing over and gene conversion is:

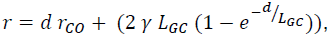

where *d* is the distance between the focal and the selected sites in bp, *r_CO_* is the rate of crossing over (/bp/female meiosis), γ is the rate of gene conversion initiation (/bp/female meiosis) and *L_GC_* is the average gene conversion tract length. Based on [31], γ = 1.25×10^−7^/bp/female meiosis and *L_GC_* =518 bp. In this study, BGS expectations were investigated using recombination rates caused by crossing over alone (*r* = *r_CO_*) and with the more realistic recombination between sites based on the combined effect of crossing over and gene conversion. When not explicitly indicated, recombination using both crossing over and gene conversion is employed.

### BGS models and parameters

We initially investigated eight BGS models using the formulas described above, with two DDFEs (log-normal or gamma; models M_LN_ and M_G_, respectively), two deleterious mutations rates (*U* = 1.2 or 0.6; models M_StdMut_ and M_LowMut_, respectively), and two recombination scenarios (with only crossovers or crossovers and gene conversion events; models M_CO_ and M_CO+GC_, respectively). For instance, the full notation of a model that uses a log-normal DDFE, a deleterious mutation rate of *U* = 0.6, and recombination that only considers crossovers is M_LN,LowMut,CO_.

To obtain the number of sites *Naa*, *Nutr_ss* and *Nnc_ss* for each 1-kb region, we followed the approach described by Charlesworth [33]. We assume 0.75 as the fraction of coding sites that alter amino acid sequences and correct this fraction by the proportion of constrained nonsynonymous sites (*cs*) to focus only on the fraction of deleterious mutations. In *D. melanogaster*, *cs* for amino acid sites is ∼0.92 [22,37] and, thus, *Naa* for a regions is *L_coding_* × 0.75 × 0.92. To obtain *Nutr_ss*, we correct the number of noncoding genic sites using the proportion of constrained sites (0.56 for introns and 0.81 for flanking UTRs [22]) and, equivalently, we use the proportion of constrained sites at intergenic sequences (∼0.5) [22,37] to obtain *Nnc_ss*.

The deleterious mutation rate per bp (*u*) will be different for different models, due to the overall mutation rate and because different DDFEs predict a different fraction of deleterious mutations with *s* < *s*_T_ that will not contribute to BGS. The two mutation rates investigated here, *U* = 0.6 (M_LowMut_) and 1.2 (M_StdMut_), represent a neutral mutation rate of 4.2×10^−9^ and 8.4×10^−9^/bp/generation. Assuming *N_e_* = 1,000,000 for *D. melanogaster*, the log-normal and gamma DDFEs described above predict 15.3 and 7.4% of deleterious mutations with *s* < *s*_T_, respectively. Therefore, the deleterious mutation rate relevant for BGS is 84.7 (M_LN_) and 92.6% (M_G_) that of the mutation rate.

#### BGS models with two DDFEs

All previous BGS models assume a single DDFE for sites under selection. As a first approximation towards more realistic description of selection across the genome, we investigated two additional models that allow for two DDFEs, one for nonsynonymous mutations and the another for deleterious noncoding mutations (each DDFE following the parameters indicated above). The *B* landscapes generated by these hybrid models are intermediate relative to the other eight models: a model assuming a log-normal DDFE for nonsynonymous mutations and a gamma for deleterious noncoding mutations (M_LN/G,StdMut,CO+GC_) shows a genome-wide median *B* of 0.469 whereas a model with a gamma DDFE for nonsynonymous mutations and a log-normal for deleterious noncoding mutations (M_G/LN,StdMut,CO+GC_) shows a median *B* of 0.459. Spearman’s rank *ρ* between estimates of *B* across the genome from these 2-DDFE models and the other eight single-DDFE models range between 0.974 and 0.998 for M_LN/G,StdMut,CO+GC_, and between 0.946 and 0.986 for M_G/LN,StdMut,CO+GC_.

#### X chromosome

For the X chromosome, we have assumed equivalent deleterious mutations rates than for autosomes and adjusted the distribution of selection coefficients to *s_X_* = 2/3 *s*_A_ (assuming equal number of breeding males and females in the population and genic selection, *h* = 0.5). Estimates of the ratio of observed neutral diversity for the X and autosomes (X/A) also assume equal number of males and females, with X/A = 0.75 × *B*_X_ / *B*_A._ Equivalently, *s*_T_ for the X chromosome varies from that for autosomes, with *s*_T_ for the X chromosome now satisfying *N_e X_ s_X_* ∼ 1. For the X chromosome, the log-normal and gamma DDFEs predict 18.6 and 9.1% of deleterious mutations with *s* < *s*_T_, respectively (both percentages higher that for autosomes, see above). Finally, we allowed for an effective difference in recombination rates due to the lack of meiotic recombination in *Drosophila* males, with 0.5*r* and 0.66*r* for autosomes and the X chromosomes, respectively. For each of the BGS models under study, we obtained specific solutions to the integrals described above along a continuum of recombination rates for the X chromosome and autosomes separately.

#### Complete and trimmed chromosomes

All analyses were carried out based on the study of complete chromosome arms as well as after removing euchromatic regions near to centromeres and telomeres with evident reduction in crossover rates [133]. Following [26] we use the term *trimmed chromosomes* or *trimmed genome* to designate genomic regions after removing sub-centromeric and -telomeric regions with strongly reduced crossover rates. Sub-centromeric regions with reduced crossover rates were assigned by starting at the centromere and moving into the chromosome arm until a minimum of 3 consecutive 100-kb windows showed crossover rates >1 cM/Mb. Sub-telomeric regions with evident reduction in crossover rates were assigned in an equivalent manner, by starting at the telomere and moving towards the center of the chromosome until a minimum of 3 consecutive 100-kb windows showed >1 cM/Mb.

### Estimates of neutral nucleotide diversity across the *D.melanogaster* genome

Nucleotide diversity across the *D. melanogaster* genome was estimated from a sub-Saharan African population (Rwanda, RG, population of the Drosophila Population Genetics Project, DPGP; www.dpgp.org/ and [62]). *D*. melanogaster is thought to have originated in sub-Saharan Africa, and eastern Africa—including Rwanda—in particular [134]. Our use of the RG population, therefore, minimizes the non-equilibrium effects caused by recent expansion observed in western Africa and non-African *D. melanogaster* populations [62,134]. Additionally, the RG population combines a relatively large sample (n=27), and low and well characterized levels of admixture [62]. We followed Pool et al. (2012) and used updated assemblies and fasta files from the DPGP2.v3 site (see http://www.dpgp.org/dpgp2/DPGP2.html and [62] for details). We analyzed these assemblies with, 1) sites putatively heterozygous or with quality value smaller than Q31 masked to ‘N’, and 2) putatively admixed regions of African genomes filtered to “N” based on the description of admixed regions from [62] (also available from http://www.dpgp.org/dpgp2/DPGP2.html). Finally, we only investigated non-N sites that were present in a minimum of 10 sequences. Equivalent results were obtained when the analyses were restricted to sites with a minimum of 15 sequences, or after removing regions showing excess admixture as well as regions showing excess long-range identity-by-descent (IBD) [62] (data not shown).

Neutral (silent) diversity was estimated as pairwise nucleotide variation per site at intergenic sequences and introns (π_sil_) and following the approach described in [31]. In short, we first annotated all gene models, transposable elements and repetitive sequences onto the reference sequence allowing for overlapping annotation. Intergenic sites correspond to sites between gene models (excluding annotated UTRs), transposable elements or repetitive sequences. Intronic sites are analyzed as such only when they never overlap with another annotation (e.g., with alternatively spliced exons or other elements within introns). After this filtering approach, intergenic and intronic sites show similar levels of silent diversity, with a very weak tendency for π_sil_ in introns (π_introns_ = 0.0085) to be higher than in intergenic (π_intergenic_ = 0.0082) sites (Sign test, *P* = 0.034 and *P* = 0.056, along autosomes and X chromosome, respectively).

Unless indicated otherwise, diversity analyses were performed using all 100-kb non-overlapping windows across the genome whereas analyses of 10-kb and 1-kb regions were limited to regions with more than 1,000 and 500 silent sites, respectively. Inferences about the frequency of segregating variants within the population were based on estimates of Tajima’s *D* [69] after normalizing by *D*min (*D*/*D*min) following Schaeffer (2002) [135]. Equivalent conclusions were obtained when using DGRP sequenced strains [136] from a North American natural population (Raleigh, NC, USA).

#### Outliers of nucleotide diversity

To parameterize departures of observed levels of diversity π_sil_ relative to the levels predicted by *B*, we applied a generalized regression model (GRM), and obtained regression residuals π_sil-R_. In particular, we obtained studentized residuals to detect outlying values and obtain associated probabilities. Studentized residuals are equivalent to standardized residuals (the ratio of the residual to its standard error) and have a Student’s *t* distribution, but in the case of studentized residuals the data point or observation under study (possibly an outlier) is removed from the analysis of the standard error. Because the sample size in our analyses is very large and the fraction of outliers small, studentized and standardized residuals are almost equivalent and generate the same outiler regions at the three physical scales analyzed. We applied GRM separately to autosomes and X chromosome to prevent including assumptions about the true X/A ratio. To correct for multiple tests, we applied the Benjamini and Hochberg (1995) method with FDR = 0.10.

### Estimates of rates of protein evolution

#### Genes and sequence alignments

We obtained CDS sequence alignments in the 5 species of the *D. melanogaster* subgroup (*D. melanogaster*, *D.simulans*, *D. sechellia*, *D. yakuba* and *D. erecta*) based on multiple-species alignments available from the UCSC Genome Browser (http://hgdownload.cse.ucsc.edu/goldenPath/dm3/multiz15way/alignments/flyBaseGene.exonNuc.fa.gz) and current gene annotation and genomic location in *D. melanogaster* (http://flybase.org; *D. melanogaster* annotation release 5.47, 10/9/2012). We removed genes with coding sequences in *D. melanogaster* shorter than 450 bp or with fewer than 100 amino acids positions present in all five species after alignment. We also removed genes presenting premature stop codons in at least one species relative to the stop codon position in *D. melanogaster*. Finally, for genes with multiple transcript forms we chose the CDS alignment corresponding to the longest transcript. In total, we investigated rates of evolution in 6,876 protein-coding genes.

#### Rates of protein evolution

We obtained *d*_N_ (the number of nonsynonymous substitutions per nonsynonymous site), *d*_S_ (the number of synonymous substitutions per synonymous site) and *ω* (the ratio *d*_N_/*d*_S_) for each gene. To this end, we used the program *codeml* as implemented in PAML v 4.5 [137,138] and applied a branch model that allowed different *ω* in all internal and external branches of the five-species tree. We focused on *ω* along the *D. melanogaster* branch after split from the *D.simulans* lineage to investigate recent patterns of efficacy of selection on amino acid changes and the possible association with BGS effects across the genome absed on recombination rates estimated in *D. melanogaster*.

We followed Larracuente et al. (2008) [84] to prevent combining genes with amino acids evolving under positive selection and genes with most amino acids evolving under a nearly neutral scenario, either purifying selection or neutrality. We, therefore, applied *codeml* and compared Model M1a (nearly neutral evolution) against a model that allows the additional presence of positive selection at a fraction of sites (model M2a). In particular, we compared maximum likelihood estimates (MLEs) under these two models and applied a likelihood ratio tests (LRTs) [139,140] to detect differences between the models. A total of 125 genes showed evidence of positive selection in the *D. melanogaster* lineage after applying the Benjamini and Hochberg (1995) method to correct for multiple tests (FDR = 0.05). To further reduce the incidence of genes under positive selection and/or recent pseudogenization we removed genes showing *ω* greater than 0.75 (note that the median *ω* in the *D. melanogaster* lineage is 0.0696). In all, we investigated 6,677 genes with no evidence of positive selection or drastic reduction in constrains along the *D. melanogaster* lineage.

Finally, we took into account the consequences of variable recombination and *B* on the efficacy of selection on synonymous mutations. In *D. melanogaster*, synonymous mutation are under weak selection ([113] and references therein) and therefore a reduction in efficacy of selection is predicted to increase *d*_S_, possibly biasing the direct use of the ratio *d*_N_/*d*_S_. On the other hand, the inclusion of *d*_S_ corrects for possible differences in coalescent times and ancestral polymorphism across the chromosome. We then applied generalized linear models, GRM, and obtained regression residuals of *ω* along the *D. melanogaster* lineage after controlling for *d*_S_ (*ω*_R_) and used *ω*_R_ as an estimate of variable efficacy of selection on amino acid changes. Equivalent results are obtained when we used regression residuals of *d*_N_ along the *D. melanogaster* lineage after controlling for *d*_S_ (*d*_N-R_), as an estimate of variable efficacy of selection on amino acid changes. We obtained regression residuals of *ω* or *d*_N_ for autosomes and the X chromosome separately.

### Forward computer simulations and estimates of the proportion of adaptive substitutions

We used the program SLIM [94] to capture the consequences of temporal changes in recombination rates on estimates of α. Simulations followed a panmictic population of 10,000 diploid individuals (*N*) and a chromosome segment of 1 Mb that contained 100 protein encoding genes evenly distributed, one every 10 kb. Each 10-kb region included a typical *Drosophila* gene: a 1,000 bp 5’ UTR, a first short 300-bp exon, a 1,000 bp first intron, two additional 600 bp exons, a short 200-bp internal intron, and a 300 bp 3’-UTR, followed by a 5,000 bp intergenic sequence. Mutations were assigned to have the same parameters than those in the BGS models described above, with two mutation types: neutral and deleterious. The population mutation rate was set to *Nu* = 0.005 and deleterious mutations were assumed to follow a gamma DDFE (*h* = 0.5) with average *Ns* = −2,500 [33,38]. The proportion of deleterious mutations at the different genomic elements was set to 0.92 at first and second codon positions, 0.81 at UTRs, 0.56 at introns, and 0.5 at intergenic sites [22,37]; all third codon positions evolved neutrally. Note that the simulation of a smaller population would prevent the study of deleterious mutations with scaled selection close to those estimated in *Drosophila* (*Ns* = −2,500) while the simulation of shorter genomic sequences would severely underestimate BGS effects under partial linkage.

Simulations followed 10 independent populations a minimum of 50 *N* generations after reaching equilibrium (>10 *N* generations). Recombination rates were uniformly distributed, with a population crossover rate (/bp) of *Nr_CO_* = 0.04 and *Nr_CO_* = 0.0025 for periods of high and low recombination, respectively. To prevent overestimating BGS effects we also included a constant rate of gene conversion initiation (/bp) of *N*γ = 0.05 and average gene conversion tract length of *L_GC_* =518 [31]. After the initial 10*N* generations, we sampled the population every 0.1*N* generations, obtained polymorphism data from 20 randomly drawn chromosomes, and compared them to a sequence that evolved independently for 20*N* generations to obtain divergence (*d*) values. These levels of polymorphism and divergence are equivalent to those observed within *D. melanogaster* for polymorphism and between *D.melanogaster-D.simulans* for divergence, where τ = *d*/*θ* = ∼6 for neutral sites [95].

Estimates of the proportion of adaptive substitutions (*α*) at selected sites was restricted to the 100-kb central region. Following Eyre-Walker and Keightley [91], α was obtained by maximum likelihood (ML) to capture the presence of strongly deleterious mutations in the simulations, with and without the possibility of population size change, and with and without correcting for the contribution of polymorphism to divergence [95]. Estimates of α were obtained using the DFE-alpha server (http://lanner.cap.ed.ac.uk/∼eang33/dfe-alpha-server.html).

## Acknowledgments

The author thanks Ana Llopart, Andrew Adrian and Brian Charlesworth for comments and suggestions on the manuscript, John Pool for making publicly available sequences from the Rwanda RG population (DPGP2), and the editor and three reviewers for their constructive comments that greatly improved the manuscript.

## SUPPORTING INFORMATION

**Figure S1. Frequency distribution of *B* estimates from BGS models that differ in the distribution of deleterious fitness effects (DDFE) and deleterious mutation rate. (A)** *B* estimates based on model M_LN,StdMut_ (log-normal DDFE) and model M_G,StdMut_ (gamma DDFE), when the diploid deleterious mutation rate is *U* = 1.2. **(B)** *B* estimates based on model M_LN,LowMut_ (log-normal DDFE) and model M_G,LowMut_ (gamma DDFE), when the diploid deleterious mutation rate is *U* = 0.6 (see text for details). All results based on the analysis of 1-kb non-overlapping regions.

**Figure S2. Distribution of silent diversity (**π**_sil_) and predicted BGS effects (*B*).** Estimates of *B* based on model M_LN,StdMut,CO+GC_. Results shown for 100-kb non-overlapping regions across chromosome arm 3L.

**Table S1.** Summary of *B* estimates for the different BGS models.

| | | Deleterious Mutation Rate ($U = 1.2$) | | | | Low Deleterious Mutation Rate ($U = 0.6$) | | | |
| --- | --- | --- | --- | --- | --- | --- | --- | --- | --- |
|  |  | Log-Normal DDFE |  | Gamma DDFE |  | Log-Normal DDFE |  | Gamma DDFE |  |
|  |  | <u>Rec. CO+GC</u> | <u>Rec. CO</u> | <u>Rec. CO+GC</u> | <u>Rec. CO</u> | <u>Rec. CO+GC</u> | <u>Rec. CO</u> | <u>Rec. CO+GC</u> | <u>Rec. CO</u> |
| | | $M_{LN,StdMut,CO+GC}^*$ | $M_{LN,StdMut,CO}$ | $M_{G,StdMut,CO+GC}$ | $M_{G,StdMut,CO}$ | $M_{LN,LowMut,CO+GC}$ | $M_{LN,LowMut,CO}$ | $M_{G,LowMut,CO+GC}$ | $M_{G,LowMut,CO}$ |
| **Whole genome** |  |  |  |  |  |  |  |  |  |
| All | Median | **0.591** | 0.470 | 0.428 | 0.337 | 0.769 | 0.686 | 0.654 | 0.581 |
|  | min, max | **0 , 0.897** | 0 , 0.886 | 0 , 0.870 | 0 , 0.855 | 0.005 , 0.947 | 0.002 , 0.942 | 0.002 , 0.933 | 0.001 , 0.925 |
|  | 90% CI | **0.005 , 0.800** | 0 , 0.756 | 0.001 , 0.707 | 0 , 0.658 | 0.074 , 0.895 | 0.022 , 0.870 | 0.039 , 0.841 | 0.011 , 0.811 |
| Autosomes | Median | **0.559** | 0.432 | 0.395 | 0.303 | 0.748 | 0.658 | 0.628 | 0.550 |
|  | min, max | **0 , 0.822** | 0 , 0.804 | 0 , 0.765 | 0 , 0.75 | 0.005 , 0.907 | 0.002 , 0.897 | 0.002 , 0.875 | 0.001 , 0.866 |
|  | 90% CI | **0.002 , 0.746** | 0 , 0.704 | 0.001 , 0.635 | 0 , 0.592 | 0.048 , 0.864 | 0.017 , 0.839 | 0.027 , 0.797 | 0.009 , 0.769 |
| X | Median | **0.736** | 0.645 | 0.603 | 0.509 | 0.859 | 0.804 | 0.777 | 0.714 |
|  | min, max | **0.025 , 0.897** | 0 , 0.886 | 0.003 , 0.870 | 0 , 0.855 | 0.160 , 0.947 | 0.012 , 0.942 | 0.056 , 0.933 | 0.004 , 0.925 |
|  | 90% CI | **0.075 , 0.862** | 0.003 , 0.833 | 0.013 , 0.810 | 0.001 , 0.771 | 0.277 , 0.929 | 0.057 , 0.913 | 0.116 , 0.900 | 0.024 , 0.878 |
| **Trimmed genome** |  |  |  |  |  |  |  |  |  |
| All | Median | **0.643** | 0.550 | 0.493 | 0.416 | 0.802 | 0.742 | 0.702 | 0.645 |
|  | min, max | **0.191 , 0.897** | 0.004 , 0.886 | 0.074 , 0.870 | 0.006 , 0.855 | 0.438 , 0.947 | 0.060 , 0.942 | 0.271 , 0.933 | 0.077 , 0.925 |
|  | 90% CI | **0.411 , 0.814** | 0.218 , 0.776 | 0.238 , 0.730 | 0.137 , 0.684 | 0.641 , 0.902 | 0.467 , 0.881 | 0.488 , 0.854 | 0.370 , 0.827 |
| Autosomes | Median | **0.619** | 0.517 | 0.467 | 0.386 | 0.787 | 0.719 | 0.683 | 0.621 |
|  | min, max | **0.191 , 0.822** | 0.004 , 0.804 | 0.074 , 0.765 | 0.006 , 0.75 | 0.438 , 0.907 | 0.060 , 0.897 | 0.271 , 0.875 | 0.077 , 0.866 |
|  | 90% CI | **0.392 , 0.757** | 0.196 , 0.719 | 0.223 , 0.654 | 0.12 , 0.615 | 0.626 , 0.870 | 0.443 , 0.848 | 0.472 , 0.809 | 0.346 , 0.784 |
| X | Median | **0.761** | 0.683 | 0.641 | 0.561 | 0.873 | 0.827 | 0.801 | 0.749 |
|  | min, max | **0.471 , 0.897** | 0.147 , 0.886 | 0.226 , 0.870 | 0.091 , 0.855 | 0.687 , 0.947 | 0.385 , 0.942 | 0.476 , 0.933 | 0.302 , 0.925 |
|  | 90% CI | **0.576 , 0.866** | 0.386 , 0.838 | 0.388 , 0.819 | 0.249 , 0.781 | 0.760 , 0.931 | 0.623 , 0.916 | 0.623 , 0.905 | 0.500 , 0.884 |

\* Model $M_{LN,StdMut,CO+GC}$ is the default model.

**Table S2.** Complete distribution of *B* estimates for the different BGS models. Table shows estimates of *B* based on eight BGS models, along all chromosome arms, and for adjacent 1-kb regions.

**Table S3.** Pairwise Spearman’s rank correlation coefficients (*ρ*) between estimates of *B* from different BGS models.

| | $M_{LN,CO+GC,StdMut}$ | $M_{LN,CO,StdMut}$ | $M_{G,CO+GC,StdMut}$ | $M_{G,CO,StdMut}$ | $M_{LN,CO+GC,LowMut}$ | $M_{LN,CO,LowMut}$ | $M_{G,CO+GC,LowMut}$ |
| --- | --- | --- | --- | --- | --- | --- | --- |
| $M_{LN,CO,StdMut}$ | 0.98784 | | | | | | |
| $M_{G,CO+GC,StdMut}$ | 0.99792 | 0.98569 | | | | | |
| $M_{G,CO,StdMut}$ | 0.99132 | 0.99717 | 0.99303 | | | | |
| $M_{LN,CO+GC,LowMut}$ | 0.99999 | 0.98760 | 0.99782 | 0.99105 | | | |
| $M_{LN,CO,LowMut}$ | 0.98798 | 0.99998 | 0.98579 | 0.99713 | 0.98775 | | |
| $M_{G,CO+GC,LowMut}$ | 0.99796 | 0.98560 | 0.99999 | 0.99293 | 0.99786 | 0.98570 | |
| $M_{G,CO,LowMut}$ | 0.99141 | 0.99715 | 0.99310 | 0.99999 | 0.99114 | 0.99714 | 0.99300 |

\* See Materials and Methods for detailed description of the eight BGS models. $P < 1 \times 10^{-12}$ in all cases.

